# gaftools: a toolkit for analyzing and manipulating pangenome alignments

**DOI:** 10.1101/2024.12.10.627813

**Authors:** Samarendra Pani, Fawaz Dabbaghie, Tobias Marschall, Arda Söylev

**Affiliations:** Heinrich Heine University Düsseldorf, Medical Faculty, Institute for Medical Biometry and Bioinformatics, Moorenstr. 5, 40225 Düsseldorf, Germany; Center for Digital Medicine, Heinrich Heine University, Moorenstr. 5, 40225 Düsseldorf, Germany

## Abstract

Linear reference genomes are ubiquitously used in genomics research, despite known biases associated with their use. In recent years, there has a been a shift towards using graph-based reference genomes to address some of these biases, which has required new algorithms and file formats to be developed. This has created the need to develop new tools capable of utilizing these formats and performing operations similar to those carried out by traditional methods.In this paper we present “gaftools”, a multi-purpose tool that introduces several utilities for processing graph alignments in GAF format. Gaftools enables users to index and sort alignments, with graph ordering serving as a necessary step for the sorting process. Additionally it allows users to view subsets of alignments and perform realignment using the wavefront alignment algorithm, among other useful features. Many of these functionalities are inspired by SAMtools, which provide similar operations for linear genomes, while gaftools adapts and extends them for pangenomes.

**Availability:** gaftools is available under MIT license at https://github.com/marschall-lab/gaftools.

## 1. Introduction

Linear reference genomes are inadequate to represent the genetic diversity of a species, which leads to several limitations, including population-specific read mapping biases (1, 5, 16). With the recent efforts of the Human Pangenome Reference Consortium (HPRC), a first draft human pangenome reference has been generated (11). In parallel with this effort, several tools have emerged for graph construction (6, 10), read alignment on graphs (3, 10, 18), variant discovery and genotyping (4, 5, 17, 21), and other graph manipulation tasks (2, 5, 8, 15).

Computationally, two key components are central to most of these tools: representing a pangenome graph and the alignments to it. In this context, the most widely accepted formats are GFA (Graphical Fragment Assembly) for graphs and GAF (Graph Alignment Format) for alignments. Although a few tools have been developed to work with GFA files, there is a notable lack of tools designed to manipulate and analyze alignment files for pangenomes, comparable to SAMtools (9), an essential tool in the linear genome domain.

Here, we present *gaftools*, a toolkit offering various features for GAF alignments, such as viewing, sorting, indexing, and realigning GAFs. It also preprocesses rGFAs, adding tags essential for viewing and sorting. It is an easy-to-install tool available as a PyPI package and on Bioconda (7).

## 2. Methods

In the following sections, we describe the main functionalities of gaftools and its different utilities.

### Indexing and viewing of GAF files

The “view” command allows filtering a GAF file to alignments to specific nodes, paths or regions. This also includes coordinate system conversion of alignment paths between “stable” and “unstable” (10). The unstable system indexes each base by the node ID (a.k.a. segment id) and the offset on the node, whereas the stable system uses contig ID, independent of the graph nodes. Thus, by using “`gaftools view -format`”, conversion between the two coordinate systems is possible. As an example, a read from NA12878 genome, aligned to “>s249732>s326404>s249733” in the HPRC-r518 T2TCHM13 Minigraph based on the unstable coordinate system can be converted to the same path of “>chr19:18591208-18685851>GRCh38#0#chr19:18550488-18550808>chr19:18685851-18688039” in stable coordinate.

We also present a basic index structure for the GAF files, which is essential for the view command to enable rapid access to alignments associated with user-specified nodes and regions. The index is an inverse dictionary lookup, where the keys represent node IDs, and the values are lists of offsets within the GAF file indicating the locations where each node ID is found. Along with a block gzip (bgzip) based compression of the GAF files, the offsets enable the view command to effectively traverse the GAF file and retrieve the desired alignments.

### GFA ordering and sorting of alignments

Efficient queries on large-scale datasets necessitate ordered and sorted data. However, in contrast to a linear reference, a general graph with cycles does not immediately imply a specific ordering. To address this, we propose an ordering and sorting approach specifically designed for pangenomes. To order pangenome graphs (using “`gaftools order_gfa`”), we leverage the properties of rGFAs generated by Minigraph. rGFAs are generated by starting with a linear graph and incorporating variants through alignments, ensuring the presence of a linear reference path within the graph that serves as a backbone for ordering. We identify all the biconnected components that represent different variants in the graph, resulting in a continuous chain of these components for each chromosome. Here, we call these biconnected components “bubbles”. The articulation points of the bubbles here correspond to a reference contig, due to the way Minigraph builds these graphs from the linear reference, and we refer to these articulation points as “scaffold nodes”.

For this purpose, we introduce additional tags to the nodes of the graph, termed BO (Bubble Order) and NO (Node Order). The BO tags sequentially label the bubbles detected in each connected component, where each component represents a chromosome. We begin at the scaffold node with the smallest reference coordinates (indicating the chromosome’s start). We then follow the detected bubbles, assigning the first scaffold node (the bubble source) and all nodes within that bubble the same BO tag, corresponding to the bubble number, incremented by 1 from the previous bubble. The NO tags sequentially label the nodes inside the bubble, starting from 0, based on the lexicographic order of the node IDs. Figure 1A shows a simple chain of 4 bubbles and 5 scaffold nodes. All nodes within a bubble are assigned the same BO tag. For the NO tag (Figure 1B), scaffold nodes are assigned a value of 0, with subsequent nodes within the bubble incrementing by 1.

**Fig. 1.**
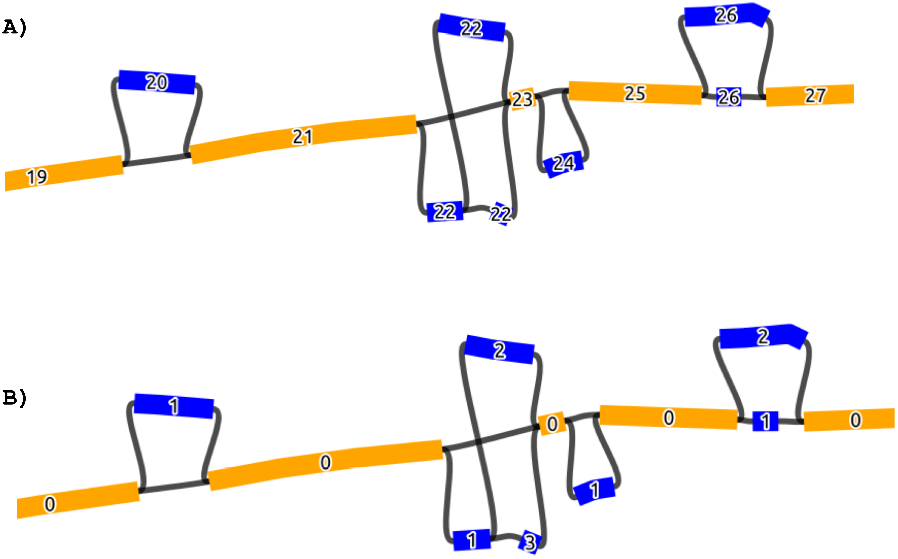
The figure depicts the BO (A) and NO (B) tags. Blue nodes are the bubble and orange ones are the scaffold nodes.

With this ordered graph structure, a sorted GAF file can be generated (“`gaftools sort`”) using the BO and NO tags as sorting keys in that order of precedence. This is necessary for various operations in order to optimize data processing and reduce redundant memory accesses.

### Exact gap-affine realignment

Sequence-to-graph alignment is a challenging task and different tools come with different limitations. GraphAligner, for instance, is based on an edit distance model, which can produce biologically implausible gap placements. With “`gaftools realign`”, we allow for affine gap-cost based realignment of each alignment to its given path using the wavefront alignment algorithm (WFA) (12, 13), which can be run in a multi-threaded mode to enhance performance. Figure 2 shows one such scenario where an alignment of NA12878 reads using GraphAligner to the HPRC pangenome obscures a deletion of 51bps but realignment of the same reads to the same positions enables the deletion to be captured.

**Fig. 2.**
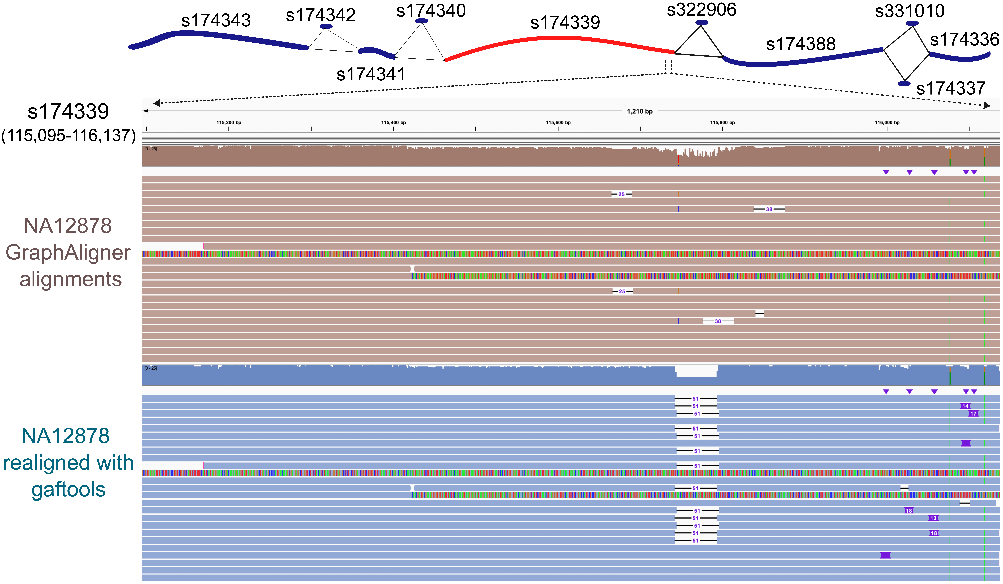
The IGV (19) figure illustrates the ONT reads of NA12878 aligned to node s174339 in the HPRC-r518 T2T-CHM13 Minigraph pangenome graph using GraphAligner (depicted in brown), compared to the same alignments after being realigned with gaftools (depicted in blue). The pangenome path is displayed above and the red node is the source of the variant.

### Additional features

gaftools also provide additional functionalities such as adding phasing information to alignments, determining the genomic path of a given node order, and generating statistics. First, phasing allows to deduce which haplotype a reads maps to in a given GAF file. Using “`gaftools phase`”, we append this information to the alignment file in the form of “ps:Z” and “ht:Z” tags, corresponding to phase set and haplotype, respectively. This data is based on “`whatshap haplotag`” TSV file output given as input (14). Additionally, we allow for retrieving the genomic sequence corresponding to a specified GFA path (e.g., “>s106519>s106520s488457>s106530>s106531”) using “`gaftools find_path`”, facilitating various downstream analysis. Furthermore, we provide comprehensive statistics for read alignments including the number of primary/secondary alignments, aligned base counts, the number of reads with at least one alignment, mapping quality, sequence identity and map ratio. In extended mode (using “`-cigar`” flag), we report additional information from the CIGAR string of the GAF file, including the total number of insertions, deletions, substitutions and match regions.

## 3. Results

We assessed the runtime and memory consumption of each gaftools command using the graph alignments of NA12878 ONT (Oxford Nanopore Technologies) reads (∼ 14X depth of coverage) aligned to HPRC-r518 T2T-CHM13 using Minigraph (Table 2). Results show that gaftools is very fast and memory-efficient for all commands, with the exception of ‘realign’. The higher runtime and memory usage for ‘realign’ are expected due to the computational demands of WFA alignment. In particular, our runtime when running on a single CPU core is comparable to Graphaligner when using 24 CPU cores, making `gaftools realign` a comparatively lightweight postprocessing step to tools like Graphaligner. We note that gaftools also supports multithreaded processing for `realign`, enabling even faster execution.

**Table 1.**
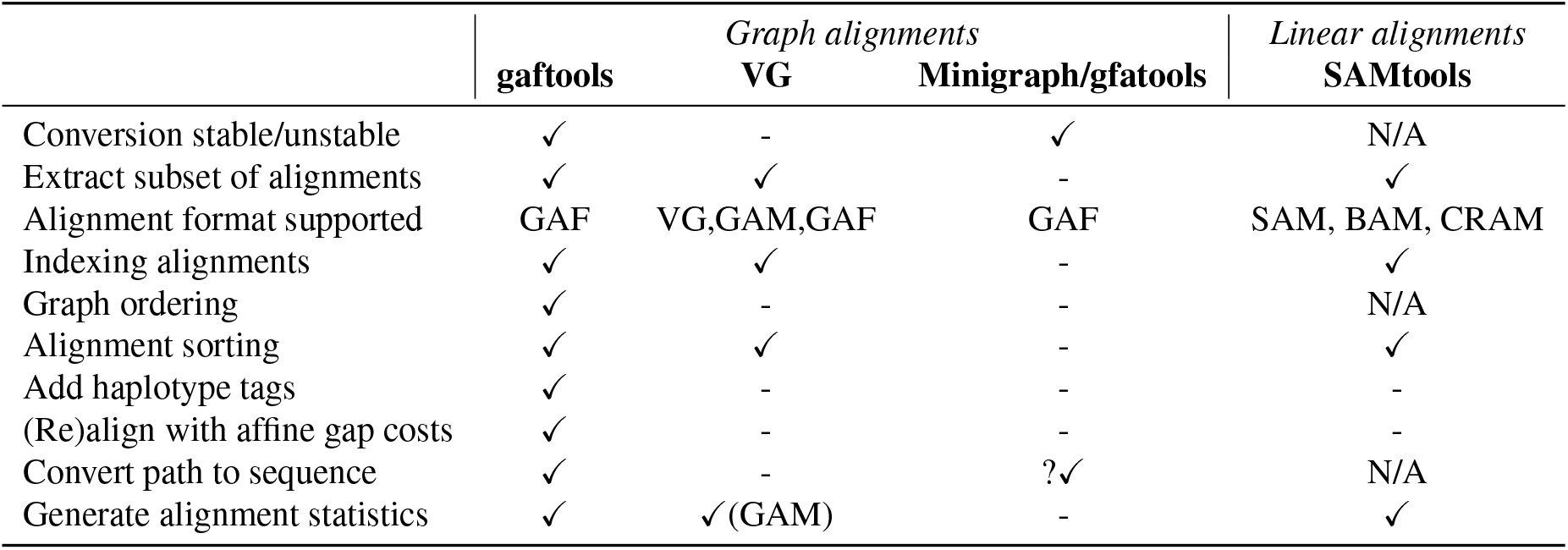
The feature matrix outlines the functionalities of gaftools, alongside other tools offering similar capabilities.

**Table 2.** Runtime and memory consumption of gaftools functionalities are given below. We performed the analysis using a single CPU core, with an Intel(R) Xeon(R) Gold 6136 CPU @ 3.00GHz with 48 CPUs and 196 GB RAM machine.

| Command | Runtime (h:mm) | Memory (GB) |
| --- | --- | --- |
| view | <0:01 | <1 |
| view (--format) | 0:20 | 2.2 |
| index | 0:01 | 2.2 |
| order_gfa | 0:01 | 2.2 |
| sort | 0:08 | 3.2 |
| phase | 0:05 | 1.8 |
| realign | 64:34 | 47 |
| find_path | <0:01 | 5.5 |
| stat | 0:04 | 1.7 |
| stat (--cigar) | 0:40 | 1.7 |

Gaftools offers key functionalities not included in state-of-the-art tools such as VG (5) and Minigraph/gfatools (8, 10) that work with GFA graphs and GAF alignments, and SAMtools (9) that works with SAM/BAM alignments (alignments against a linear reference). Table 1 shows the shared and unique features, showing that gaftools offers functionalities previously only available for linear alignments. We would also like to note that for the stable/unstable conversion, Minigraph is able to produce alignments with either stable or unstable coordinates, however, the format need to be specified before running the alignment, i.e. we would need to run the alignment step again if we wanted to change the format.

## 4. Discussion

While the tool ecosystem to work with pangenomes is maturing, there are still considerable gaps compared to software infrastructure available to work with linear references. Here, we introduced gaftools as a versatile toolkit to support the practical use of pangenome references and provide missing functionalities. In particular, gaftools focuses on working with alignments of reads to pangenome reference graphs and offers a range of functionalities for the manipulation and analysis of GAFs, including viewing alignments, sorting, and indexing, which can be viewed as counterparts to common operations on linear alignments stored in SAM/BAM files. Beyond this, gaftools offers further functionalities like coordinate conversion between the stable and unstable coordinate systems, gap-affine realignment, and transferring of phase information into GAFs. gaftools has already proven valuable for the analysis for the ONT-1KG data set (20), and we envision it will be instrumental for implementing future work-flows for pangenome-based analyses.

## Data availability

HPRC T2T-CHM13 pangenome graph can be found at https://zenodo.org/record/6983934. NA12878 GAF file can be found in https://ftp.1000genomes.ebi.ac.uk/vol1/ftp/data_collections/1KG_ONT_VIENNA/gaf/

